# Live imaging and multimodal profiling reveal transdifferentiation of a cochlear supporting cell subpopulation upon Notch inhibition

**DOI:** 10.1101/2025.11.04.686551

**Authors:** Lama Khalaily, Shahar Kasirer, Katherine Domb, Mi Zhou, Buwei Shao, Shahar Taiber, Ran Elkon, Litao Tao, David Sprinzak, Karen B. Avraham

## Abstract

Regeneration enables organisms to repair damaged tissues, yet this capacity is strikingly limited in the cochlear sensory epithelium, essential for sound detection. A major cause of hearing loss arises from the irreversible loss of sensory hair cells (HCs) in the cochlea. While supporting cells (SCs) have a latent ability to transdifferentiate into HCs, this regenerative potential is rapidly lost after development. Using live imaging and single-cell multi-omics of cochlear explants, we uncovered the cellular and molecular heterogeneity underlying the limited regenerative capacity of the neonatal mouse cochlea. Notch repression broadly silenced key SC genes, yet only a rare subpopulation of Deiters cells (DC), termed transdifferentiating DCs (tDCs), initiated the transdifferentiation into HC fate. These cells underwent coordinated transcriptional and enhancer remodeling, linking epigenetic priming with morphological plasticity, while other SCs remained refractory despite robust Notch targets downregulation. Our study provides a molecular definition of an early induced transitional DC to HC state, revealing Notch inhibition as a selective trigger that unmasks rare regenerative competence.

## Introduction

Regeneration is a fundamental biological process that allows organisms to repair and replace damaged tissues or lost structures, yet its extent varies widely across different species. While amphibians and fish can rapidly regenerate entire organs or body structures, mammals have a more restricted regeneration capacity, largely limited to tissues such as the liver, blood, and skin (*1–3*). This can be achieved through stem cell-driven regeneration (*4, 5*), where undifferentiated progenitor cells replenish lost tissues, or through cellular plasticity, in which differentiated cells directly transdifferentiate into new cell types (*6*). Notably, regenerative capacity is especially restricted in mammalian nervous and sensory systems, partly due to epigenetic constraints. Highly regenerative species maintain an open chromatin state that promotes cellular plasticity, whereas mammals have a more restrictive chromatin environment that impedes cellular reprogramming, especially in neural and sensory tissues.

One such tissue with severely restricted regenerative ability is the cochlear sensory epithelium, responsible for sound detection. Sensory hair cells (HCs) within the cochlea are particularly vulnerable to irreversible damage, leading to permanent hearing loss. The loss of HCs due to aging, noise exposure, ototoxic drugs, or diseases results in permanent hearing impairment, affecting millions worldwide (*7*). Despite advances in regenerative medicine, no effective regenerative therapies currently exist to restore auditory function, underscoring the urgent need to understand the molecular mechanisms governing cellular plasticity in the cochlea to develop new therapeutic approaches. The sensory epithelium of the mammalian cochlea, the organ of Corti, is a highly specialized structure composed of distinct cell types (*8*). It contains a single row of inner hair cells (IHCs) and three rows of outer hair cells (OHCs). These HCs are interspersed with specialized supporting cells (SCs), including Deiters cells (DC1-3), pillar cells (PCs), and inner phalangeal cells (IPhCs) (*8*).

The Notch signaling pathway is a key regulator of HC differentiation and regeneration, playing a crucial role in determining the fate of pro-sensory cells during cochlear development. Through lateral inhibition, Notch directs the differentiation of pro-sensory cells into either sensory HCs or non-sensory SCs, ensuring the proper cellular composition of the cochlea (*9, 10*). At the molecular level, Notch activation in SCs leads to the cleavage of the Notch intracellular domain (NICD), which translocates to the nucleus, forms an activation complex with the transcription factor RBPJ, and triggers the expression of downstream target genes (*11, 12*). Beyond development, Notch also governs the regenerative potential of the cochlea, as its inhibition in neonatal mice induces SC-to-HC transdifferentiation (*9, 13*). However, this regenerative potential is sharply restricted to early postnatal stages (*14–16*). By approximately one week of age, SCs lose their ability to transdifferentiate into HCs, suggesting that additional regulatory mechanisms, possibly epigenetic barriers or changes in signaling dynamics, contribute to this loss of plasticity (*16, 17*).

Recent studies have revealed how SCs lose their plasticity and transdifferentiation potential with age, driven by changes in both transcriptional and epigenetic patterns. Histone modifications such as H3K27 deacetylation and trimethylation silence HC genes in SCs, while H3K4me3 and H3K4me1 maintain a “poised” state for HC transdifferentiation (*16, 17*). Upon Notch inhibition, SCs can transdifferentiate into HCs, marked by a loss of H3K27me3 and an increase in H3K27ac (*16*). Additionally, DNA methylation at HC gene promoters in SCs, particularly at Atoh1 binding sites, prevents gene activation and limits transdifferentiation potential (*17*). Studies also show that H3K4me3 modifications in Lgr5^+^ progenitor cells may lead to HC regeneration after injury (*18*). These findings suggest that manipulating epigenetic pathways could offer therapeutic potential for inner ear regeneration.

Notch signaling plays a crucial role in cochlear development, yet its involvement in orchestrating SC-to-HC transdifferentiation remains poorly understood. Specifically, it remains unclear how Notch inhibition impacts the development of individual SCs and HCs, both at the transcriptomic level and in terms of cellular patterning. Therefore, it is vital to investigate this process at the dynamic, molecular and epigenetic levels, using high-resolution cellular analysis and dynamic imaging techniques, to provide insights into the cellular mechanisms and functional dynamics underlying this transition, particularly given the rapid loss of regenerative potential after birth.

Here, we sought to uncover the cellular and molecular determinants that allow regenerative competence within the cochlear sensory epithelium. Notch inhibition has previously been assumed to drive the formation of extra HCs exclusively through SC-to-HC transdifferentiation, yet the cellular origin of these additional HCs has never been directly observed. Prior studies have demonstrated that neonatal SCs retain a transient capacity to regenerate HC upon Notch inhibition in the neonatal cochlea and after HC ablation (*13, 19, 20*). The cell of origin of regenerated HCs has been demonstrated by fate mapping (*15, 21*). Using high-resolution live imaging, we demonstrate that Notch inhibition induces only a limited number of true transdifferentiation events, while a substantial fraction of the apparent “extra” HCs arise from pre-existing HCs that rearrange and reposition within the epithelium. Combining high-resolution live imaging with single-cell multi-omic profiling to simultaneously capture the transcriptional dynamics, chromatin accessibility landscapes, and spatial behaviors of individual SCs during early transdifferentiation revealed a previously unrecognized transdifferentiation-competent subpopulation of DCs that we termed transdifferentiating DCs (tDCs). The tDCs uniquely respond to Notch inhibition, initiating a coordinated program of morphological remodeling, enhancer activation and selective promoter accessibility changes at key HC regulators. By defining the molecular signatures and regulatory architecture that distinguish the tDCs from neighboring non-responsive SCs, our study provides a mechanistic framework for understanding the intrinsic heterogeneity that limits cochlear regeneration and identifies new targets for strategies to enhance HC regeneration.

## Results

### Notch inhibition reveals SC morphological plasticity and limited transdifferentiation

To investigate the impact of Notch signaling on SC transdifferentiation, we inhibited Notch signaling in cochlear explants at two developmental time points: postnatal day 0-1 (P0-1), when SCs retain regenerative potential, and postnatal day 6 (P6), when this capacity is largely lost (*16*) (Fig. 1, A to D). Cochlear explants were derived from transgenic mice expressing an Lfng-EGFP reporter, in which SCs are fluorescently labeled and EGFP reflects SC identity (Fig. 1B) (*20*). Explants were cultured and treated with the γ-secretase inhibitor compound E, preventing NICD cleavage and thus inhibiting Notch signaling (*22–24*) or with DMSO as a control for 24 h prior to analysis (see Materials and Methods) (Fig. 1C). Consistent with prior reports, at P0-1, Notch inhibition induced an apparent marked increase in Myo7a⁺ HCs, with some retaining EGFP expression, suggestive of active SC-to-HC transdifferentiation (Fig. 1C) (*9, 14, 16*). In addition, Notch inhibition caused reorganization of the sensory epithelium, resulting in altered tissue architecture at neonatal stages compared to control. Bulk RNA sequencing of sorted Lfng⁺ SCs following treatment with compound E revealed robust transcriptional responses to Notch inhibition, with a strong effect observed at P0-P1 (fig. S1A). Several canonical Notch target genes including *Hey1*, *Heyl*, *Hes1*, *Hes5*, and *Lfng* were significantly downregulated upon compound E treatment, confirming pathway inhibition (fig. S1A and S1B). In contrast, HC fate determinants such as *Atoh1* and *Hes6* were significantly upregulated (fig. S1B) at P0-1. These findings reinforce that compound E promotes transcriptional reprogramming from SCs toward a HC-like state. In contrast, P6 explants showed no comparable increase in HC numbers following Notch inhibition and the transcriptional response was markedly attenuated: Notch target genes were only modestly downregulated (Fig. 1D, fig. S1A and S1B), supporting a developmental restriction in regenerative competence (*16*). This HC increase at P0-1 was accompanied by ectopic rows of HCs, particularly in the outer hair cell (OHC) region of the apex, disrupting the normal checkerboard-like sensory epithelium pattern (Fig. 1E). Subtype-specific HC counting analysis at neonatal stages (P0-1) using a Vglut3 marker revealed a significant increase in Vglut3⁻ OHCs, with no change in Vglut3⁺ inner hair cells (IHCs) (Fig. 1E to F). Moreover, the analysis suggested that SC-to-HC fate conversion depends on the position along the base-to-apex axis of the cochlea with relatively strong effect at the apex, lower (but significant) effect in the mid turns, and no effect in the basal turn. These results strongly suggest that only a selected population of SCs, predominantly from the OHC compartment, undergo a SC-to-HC transition. These observations also align with prior studies (*16*).

**Fig. 1.**
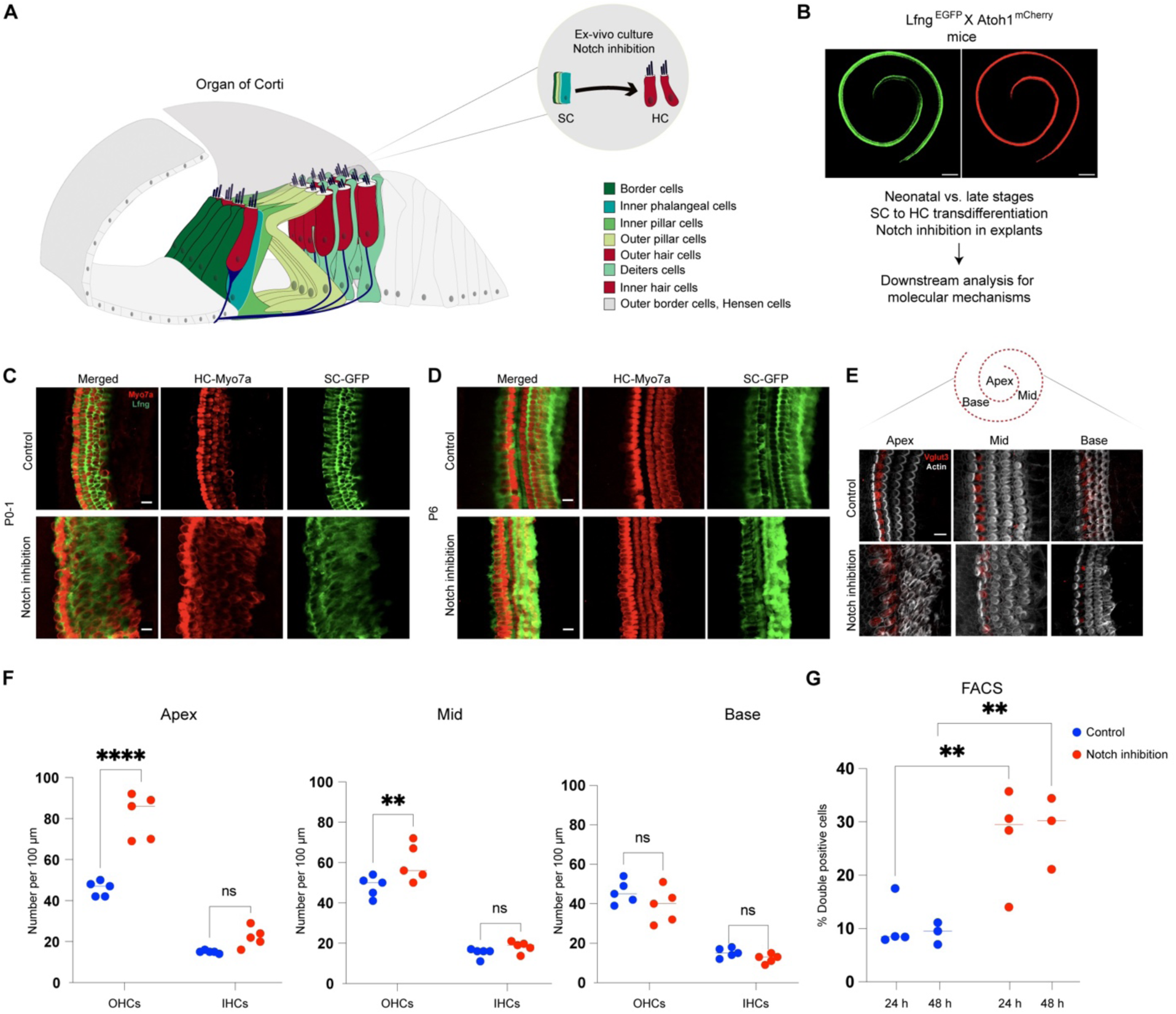
Notch inhibition induces limited SC-to-HC transdifferentiation in neonatal cochlear explants. (**A**) An illustration of the organ of Corti. Sensory inner and outer HC are labelled in red and different types of SCs labeled green. (**B**) Whole-mount immunofluorescence of P0-1 organ of Corti from Lfng^EGFP^;Atoh1^mCherry^ mice. EGFP is expressed in SCs (left) while HCs are labeled in red across the cochlea (right). (**C**, **D**) The ability of SCs (Lfng, green) to transdifferentiate into HC (Myo7a, red) in response to Notch inhibition diminishes with age. Cochleae from Lfng^EGFP^ mice at P0-1 (**C**) and P6 (**D**) were cultured and treated with compound E (2 μM) or DMSO (control) for 24 h. P0-1 samples, but not P6, show an increase in the number of HCs. (**E**) Whole-mount immunofluorescence of P0-1 organ of Corti from wild-type mice. Actin (white) marks cellular structures, and the IHC-specific marker anti-Vglut3 is shown in red. (**F**) Quantification of OHCs and IHCs at neonatal stages by immunostaining against Vglut3 across apex, mid, and base following treatment with Notch inhibitor. (**G**) FACS sorting of double-positive SCs (responsive with GFP^+^mCherry^+^) versus single-positive SCs (non-responsive with GFP^+^) upon Notch inhibition. The graph quantifies the percentage of responsive SCs, calculated as 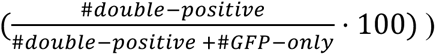. Scale bars: 100 μm (whole cochlea), 10 μm (cross sections). Data represent at least three independent biological replicates for immunostaining (C-D). Quantifications are based on n=5 biological replicates per condition for counting (F) and n=3-4 per condition for FACS (G). Statistical significance was determined using one-way ANOVA with Holm-Sidak post hoc test (**P<0.01, **** P<0.0001).

We next sought to determine whether the observed increase in HCs upon Notch inhibition originated exclusively from SC-to-HC transdifferentiation. To confirm this, we used cochlear explants from double-transgenic mice expressing both the Lfng-EGFP reporter, which marks SCs (EGFP⁺), and the Atoh1-mCherry reporter, which marks HCs (mCherry⁺) (*25*) (Fig. 1B). We then used fluorescence activated cell sorting (FACS) analysis of cochleae after Notch inhibition to determine the fraction of cells that converted from SCs to HCs. To calculate the fraction of SCs undergoing SC-to-HC transdifferentiation we divided the number of EGFP⁺mCherry⁺ double-positive cells (transdifferentiated SCs) by the number of EGFP⁺ cells (non-transdifferentiated SCs). This quantification-based approach controls for potential confounding factors such as dissociation efficiency, organ number, and cell viability. Quantification revealed a significant increase in double positive cells at both 24 and 48 h compared to controls (Fig. 1G, fig. S1C and fig. S1D), confirming a limited yet measurable transdifferentiation response. Most SCs remained EGFP⁺mCherry⁻, highlighting intrinsic heterogeneity in responsiveness to Notch inhibition.

### Live imaging reveals dynamic behavior and spatial origins of transdifferentiating SCs

While FACS analysis confirmed a limited SC-to-HC fate conversion following Notch inhibition, it could not resolve the temporal sequence or dynamic behaviors associated with this process. To date, most studies investigating Notch inhibition in the cochlea have relied on endpoint immunostaining, which captures only static snapshots of transdifferentiation and does not account for dynamic patterning changes. As a result, the real**-**time cellular dynamics including the origin, timing, and migratory behavior of responsive cells remain poorly characterized. To gain further insight, we performed live imaging on neonatal cochlear explants from Lfng^EGFP^;Atoh1^mCherry^ mice and tracked the response to Notch inhibition by compound E. Time-lapse imaging revealed that Notch inhibition caused two distinct processes: transdifferentiation from SC to HC and reorganization of cells within the OHC region. Transdifferentiation of SC to HC is observed through the initial appearance of EGFP^+^mCherry^+^ cells in the lateral side of the OHC region, beginning around 20 h post-treatment, with substantial increases by 30-48 h, particularly in the apex region (circled cells in Fig. 2A and movie S1). Such transdifferentiations were not observed in the base region or in control explants treated with DMSO (Fig. 2A, fig. S2, movie S2 and movie S3).

**Fig. 2.**
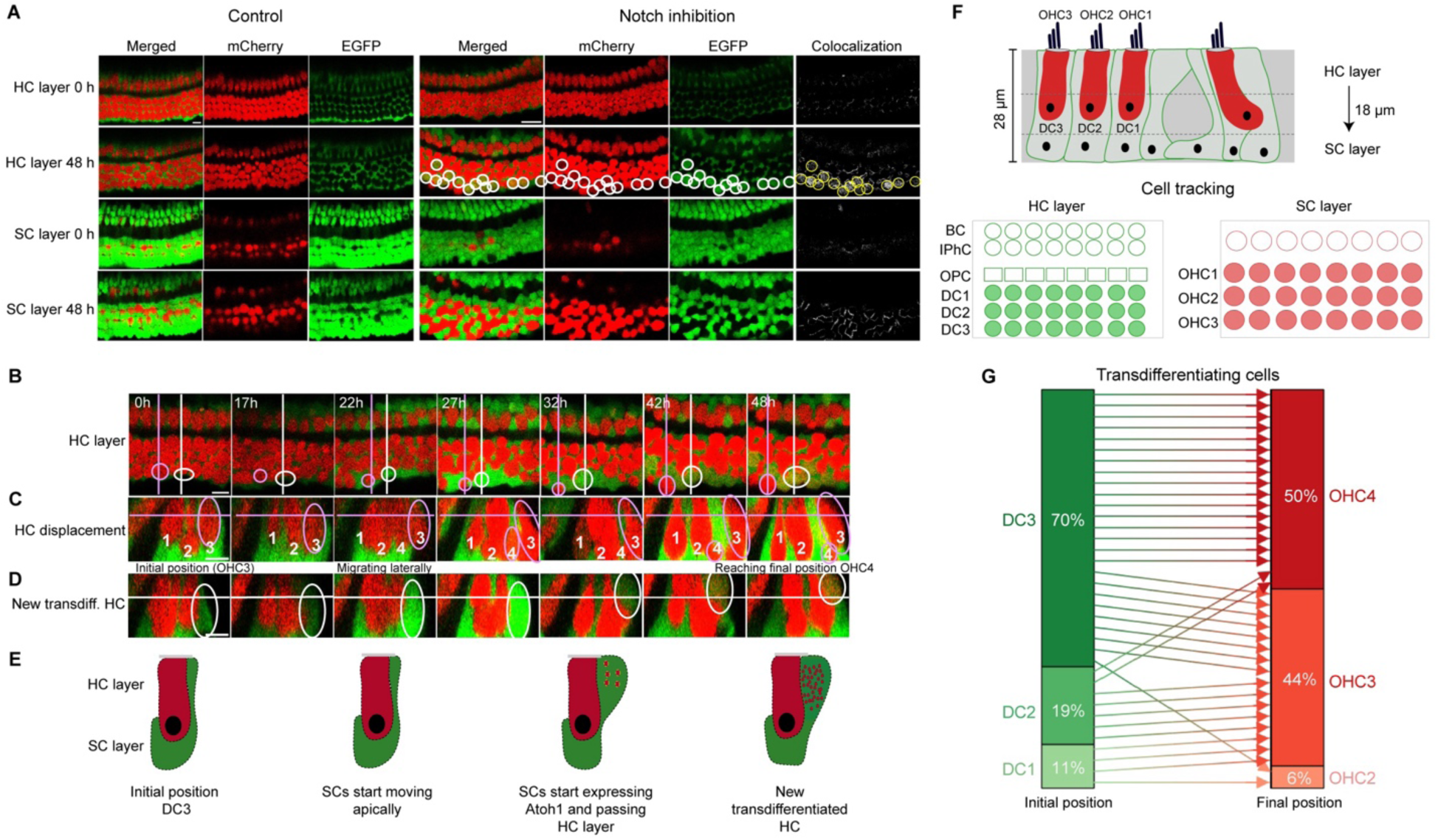
Live imaging reveals dynamic behavior and spatial origin of transdifferentiating SC following Notch inhibition. (A) Filmstrips from Lfng^EGFP^;Atoh1^mCherry^ cochlear explants apex control (DMSO) and apex treated with compound E for 48 h show the appearance of mCherry⁺ cells emerging outside the native HC rows (circles) in the apex region, whereas the control maintains precise alignment and organization of HC and SC layers. White circles highlight EGFP⁺mCherry⁺ double-positive cells, indicating transdifferentiation events. Scale bar: 10 µm. (**B-D**) Representative examples for tracking of two cells. One rearranged HC started as mCherry+ at OHC3 and ended as mCherry+ at OHC4, marked with a dashed magenta circle, and one transdifferentiating cell started as EGFP+ at DC3 and ended as EGFP+mCherry+ at OHC4, marked with a solid white circle. A top view of HC layer with both tracked cells is shown, with dashed and solid lines marking side views sections (**B**). In addition, side views of both HC displacement (**C**), each HC is labeled with consistent number according to tracking and transdifferentiating cells (**D**) are shown, with lines marking HC layer (top view section). (**E**) Schematic illustration of apical migration and marker expression of transdifferentiating cells. (**F**) Upper image: Live imaging planes used to distinguish HC and SC layers in cochlear explants. Lower image: Initial SC rows (e.g., DC1-3) and final HC rows (e.g., OHC1-4) positions. (**G**) Quantification of the initial and final rows for each tracked transdifferentiating cell across 48 h of live imaging. Each arrow represents the trajectory of an individual cell. n=4 biological repeats (movies). Total number of tracked double positive cells - 36 (6 from first movie, 18 from second movie, 4 from third movie, 8 from fourth movie).

In addition to the transdifferentiating events, we also observe significant migratory movements within the OHC region. Transdifferentiating cells originally located in the SC layer migrated apically toward the HC layer, while some HCs appeared to migrate basally towards the SC layer (Fig. 2A). These opposing, directional displacements were confined to the OHC region and were not observed on the IHC side, suggesting region-specific responsiveness to Notch inhibition.

To obtain a more detailed understanding of the transdifferentiation and migratory dynamics we tracked single cells in 3D (Fig. 2, B to E). First, we observed pre-existing HCs rearranged laterally from the third OHC layer (OHC3) to an ectopically formed fourth OHC row (OHC4) (dashed magenta circle, Fig. 2, B to C). Second, we tracked a transdifferentiating SCs emerging from the third DC row (DC3) that starts migrates apically and starts expressing mCherry, reflecting dim *Atoh1* expression (white circle, Fig. 2B, D and E). Together, these complementary trajectories demonstrate that the OHC4 row is not only the product of new HCs formed by transdifferentiation but also includes pre-existing HCs displaced due to tissue-level rearrangements. Thus, the mis-patterning of the sensory epithelium arises from the combined contributions of migration-driven repositioning and *de novo* HC generation.

To determine the specific origins and destinations of transdifferentiated HCs, we tracked individual EGFP⁺mCherry⁺ double-positive cells throughout the 48-h imaging period. Each cell’s initial identity was mapped to a defined SC subtype was mapped to a defined SC subtype DC1-3, pillar cells (PCs), inner phalangeal cells (IPhCs), or border cells (BCs) based on spatial positioning, and its final location was recorded within the HC layer at 48 h post-treatment (Fig. 2F and table S2). This analysis revealed that most transdifferentiating cells arose from DC3 (70%), with smaller contributions from DC2 (19%) and DC1 (11%) (Fig. 2G). In final destinations, about half of these cells integrated into the newly formed OHC4 row (50%), while 44% localized into the OHC3 row and a small fraction (6%) positioned into the OHC2 row. Notably, the disproportionate contribution of DC3 to OHC4 emphasizes its predominant role as the origin of *de novo* HCs.

Together, these findings reveal that Notch inhibition induces a limited SC-to-HC transdifferentiation and rearrangement process in the apex OHC region in newborn mice. Transdifferentiating cells predominantly originating from specific DCs, mostly from DC3. These cells together with migratory HCs, contribute to the formation of ectopic OHC row. Crucially, our live imaging data provide the first dynamic visualization of this process, uncovering the essential sequential trajectories leading to patterning mis-organization in response to Notch inhibition.

### Notch inhibition unmasks tDCs with HC transcriptional propensity

Previous studies have shown that inhibition of Notch signaling using γ-secretase inhibitor promotes transdifferentiation of a subset of SCs into HCs, while others remain unresponsive (*16*). Consistent with these findings, our live imaging revealed that Notch inhibition induces transdifferentiation predominantly in a specific subset of DC, while other SCs remain largely unresponsive (Fig. 2). This selective responsiveness prompts a fundamental question: What are the molecular and epigenetic features that uniquely prime DCs for HC fate conversion, and what underlies the resistance in other SC types? Here, we specifically set out to examine the earliest cellular and molecular responses to Notch inhibition, reasoning that these early events would provide direct insight into the primary effectors of fate conversion.

To address this question and investigate the molecular basis of this variability, we performed single-cell multi-omic analysis, enabling the simultaneous profiling of transcriptional activity and chromatin accessibility at single-cell resolution during the early stages of transdifferentiation. We cultured cochlear explants from P0-P1 mice in the presence of compound E or DMSO for 24 h, followed by tissue dissociation and 10x multiome library preparation to assess both RNA expression by single cell RNA-seq and chromatin accessibility by single cell ATAC-seq from the same individual cells (Fig. 3A). This approach allowed simultaneous high-resolution analysis of gene expression and chromatin accessibility within individual SCs, enabling dissection of subtype-specific responses at early stages of fate conversion.

**Fig. 3.**
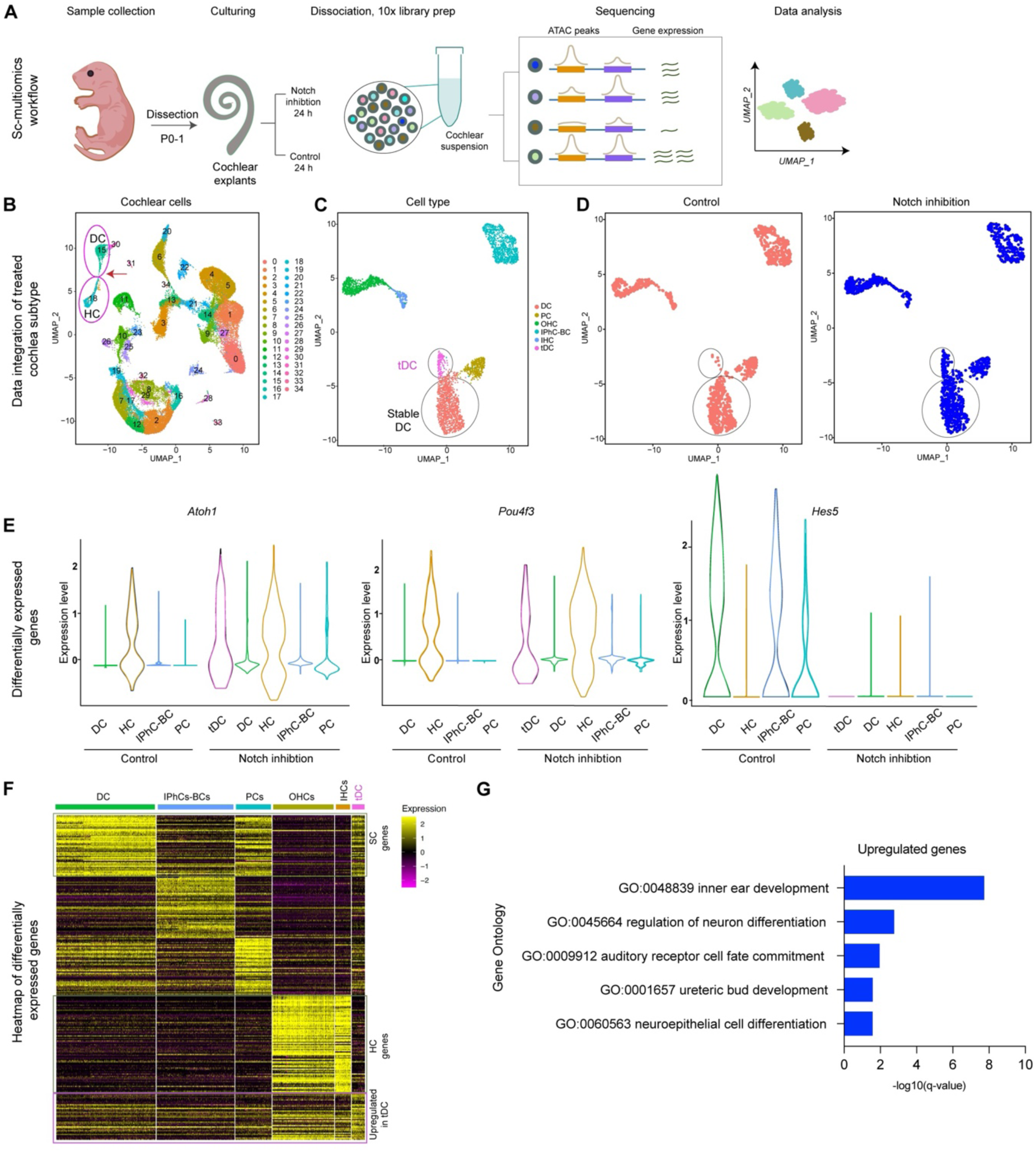
Single-cell multi-omic profiling reveals a subpopulation of SC responses to Notch inhibition and early transcriptional changes during HC fate commitment. (**A**) Schematic figure of the experimental workflow. Cochlear explants from P0-P1 mice were cultured for 24 h in the presence of Notch inhibitor (compound E) or DMSO control. Tissues were dissociated and subjected to 10x Genomics multiome profiling for simultaneous single-cell RNA expression and chromatin accessibility analysis (see also Materials and Methods section) (**B**) UMAP projection showing integrated transcriptomic clustering of cochlear cell subtypes across conditions. For main cluster ID, 18=HC, 19=IPhC-BC, 15=DC, 30=PC. Integration of control and Notch-inhibited datasets revealed marked transcriptional remodeling specifically within the DC cluster. A subset of DC acquired an intermediate transcriptional profile enriched in HC-associated genes (red arrow), forming a distinct cluster adjacent to the HC cluster. (**C** and **D**) Sub-clustering population revealed two distinct transcriptional states: black circle of “Stable DCs” and pink circle of “tDCs” based on cell type (C) and treatment (D). (**E**) Violin plots of marker gene expression: *Atoh1* and *Pou4f3* are induced in tDCs after Notch inhibition, whereas *Hes5* is downregulated in all SCs upon Notch inhibition. (**F**) Heatmap of marker genes, upregulated in each cluster compared to other clusters. Differentially expressed genes in tDCs (pink) versus stable DCs (green) highlight upregulation of HC lineage programs. (**G**) Gene ontology enrichment analysis (using Metascape online resource) of genes up regulated in tDCs compared with control DCs reveals significant activation of biological process GO terms involved in inner ear development, neuron differentiation and others. Data represent two independent biological replicates per condition.

Integration of transcriptomic data from all conditions revealed well-defined clusters corresponding to known cochlear cell types, including HCs, DCs, PCs, and IPhCs-BCs) (Fig. 3B). To validate cluster annotation, we examined the expression of canonical marker genes across the UMAP projection (fig. S3A). HC clusters were marked by *Myo7a*, *Gfi1*, and *Pou4f3*, DC/PC clusters by *Prox1* and *Fgfr3*, and IPhC-BC clusters by *Sox2*, consistent with their known molecular identities (Fig. 3, A and B) (*26*).

Integration of control and Notch-inhibited datasets revealed marked transcriptional remodeling specifically within the DC cluster. A subset of DC acquired an intermediate transcriptional profile enriched in HC-associated genes suggesting a population shift from DC towards HC (Fig. 3B and fig. S3A).

To identify responsive subpopulations within the SC population, we focused on the major sensory epithelial populations, DCs, PCs, and IPhCs-BCs, and performed additional sub-clustering. This analysis revealed two transcriptionally distinct DC states that exhibited distinct response to compound E treatment: “tDCs (cluster tDC, magenta dots) and “Stable DCs” (salmon dots) (Fig. 3C). tDCs were specifically enriched in compound E-treated samples (Fig. 3D and fig. S3C), indicating selective competence to engage in reprogramming, whereas stable DCs failed to activate HC gene programs despite uniform Notch inhibition. Marker gene projections onto the subcluster UMAP validated these assignments (fig. S3B): tDCs expressed early HC markers including *Myo7a*, *Pou4f3*, and *Gfi1*, while stable DCs retained SC identity genes such as *Sox2*, *Prox1*, and *Fgfr3*. Other SC types were identified through specific markers: PCs expressed *Sox2*, *Fgfr3*, *Prox1*, while IPhCs-BCs were characterized by *Sox2*, *Matn4*, *Fabp7*, and *Rorb* (fig. S3B). Notably, tDCs displayed a mixed transcriptional profile, characterized by induction of early HC markers (*Myo7a, Pou4f3, Gfi1*) together with retention of selected SC markers such as Sox2, while canonical DC markers including *Fgfr3* and *Prox1* were reduced (Fig. S3B). This combination supports the interpretation that tDCs represent a transitional DC-to-HC state rather than fully differentiated HCs.

Supporting this, violin plots showed that *Atoh1*, *Pou4f3*, and *Hes6* were highly expressed in tDCs when treated with compound E but not in control DMSO treatment (Fig. 3E). In contrast, stable DCs as well as PCs and IPhC/BC, did not exhibit such a response of HC specific targets to compound E. Notch effectors (*Hes1*, *Hes5*, *Heyl*) were sharply downregulated in almost all SC types following Notch inhibition (Fig. 3E and fig. S4A). While **Atoh1** activation represents an early commitment step, late HC markers such as *Rasd2* remained largely unexpressed after 24 h of Notch inhibition (fig. S4A).

Differential gene expression analysis revealed a striking transcriptional divergence between tDCs (pink) and other SC types, including stable DCs (green), PCs, and IPhCs-BCs (Fig. 3F). tDCs displayed broad downregulation of canonical SC identity genes (Fig. 3F, red box), consistent with loss of SC fate. At the same time, they exhibited partial activation of HC-associated gene programs (Fig. 3F, yellow box), reflecting early initiation of the HC lineage. In addition, tDCs displayed a unique set of genes selectively upregulated compared to both stable DCs and native HCs (Fig. 3F, purple box), suggesting the emergence of an intermediate transcriptional state rather than a complete switch. Together, this analysis highlights that tDCs do not simply silence SC programs but instead undergo a coordinated transcriptional reprogramming that combines repression of SC identity, partial activation of HC determinants, and induction of tDCs-specific genes, marking them as a distinct fate-competent subpopulation.

Gene Ontology enrichment of the tDC-upregulated gene set, identified from tDCs compound E compared to DC-DMSO, revealed a strong enrichment for developmental processes associated with sensory and neuronal differentiation (Fig. 3G and table S3). The most significantly enriched categories included inner ear development, regulation of neuron differentiation, and neuroepithelial cell differentiation. Consistent with these functions, several genes linked to cell-cell adhesion and junctional remodeling (table S3) and cell migration and cytoskeletal reorganization (*Ccbe1*, *Fgf8*, *Pak3*, *Anks1*) were upregulated, suggesting that tDCs not only activate developmental fate programs but also acquire structural plasticity required for epithelial reorganization.

Conversely, tDC-downregulated genes were enriched for processes involved in cell adhesion, migration, and junction maintenance (fig. S4 and table S3). This included repression of multiple adhesion-related genes and migration regulators (table S3), indicating a progressive loss of mature SC features. Together, these patterns imply that Notch inhibition triggers a dual remodeling process in tDCs activation of HC associated developmental programs alongside dismantling of their native adhesion and junctional architecture, enabling their transition toward a new sensory fate.

Together, these results demonstrate that Notch inhibition elicits a broad downregulation of the SC key genes across nearly all SC types, yet only a small subset of DCs, the “tDCs” can subsequently activate HC gene networks.

### tDCs undergo chromatin remodeling to adopt a HC-like fate

Having identified a transcriptionally distinct population of tDCs that emerges specifically after Notch inhibition (Fig. 3), we next sought to determine whether their competence to activate HC gene programs is linked to a distinct chromatin landscape. Because transcriptional reprogramming requires access to lineage-specific regulatory elements, we reasoned that tDCs might possess an altered epigenomic state that distinguishes them from the stable DCs and other SCs.

Global profiling of open-chromatin state revealed that tDCs diverge epi-genomically from all SC subtypes. Principal component analysis (PCA) of pseudobulk profiles showed that tDCs separated from both control DCs and other SC types along the primary variance axis, reflecting a shift of tDCs from DC/PC toward an HC-like state in both the transcriptome (sc-RNA, table S6) and in chromatin accessibility (sc-ATAC, GEO; accession number in process) (Fig. 4A). This coordinated shift in IPhC-BCs or PCs remained largely unchanged, clustering closely with their respective controls in both the transcriptomic and chromatin accessibility. The parallel separation of tDCs in both data layers demonstrates that transcriptional reprogramming in these cells is accompanied by broad remodeling of chromatin accessibility. Consistent with this, unsupervised clustering of scATAC-seq data alone resolved major cochlear cell types, and gene-level accessibility patterns closely correlated with corresponding transcriptional profiles (fig. S5).

**Fig. 4.**
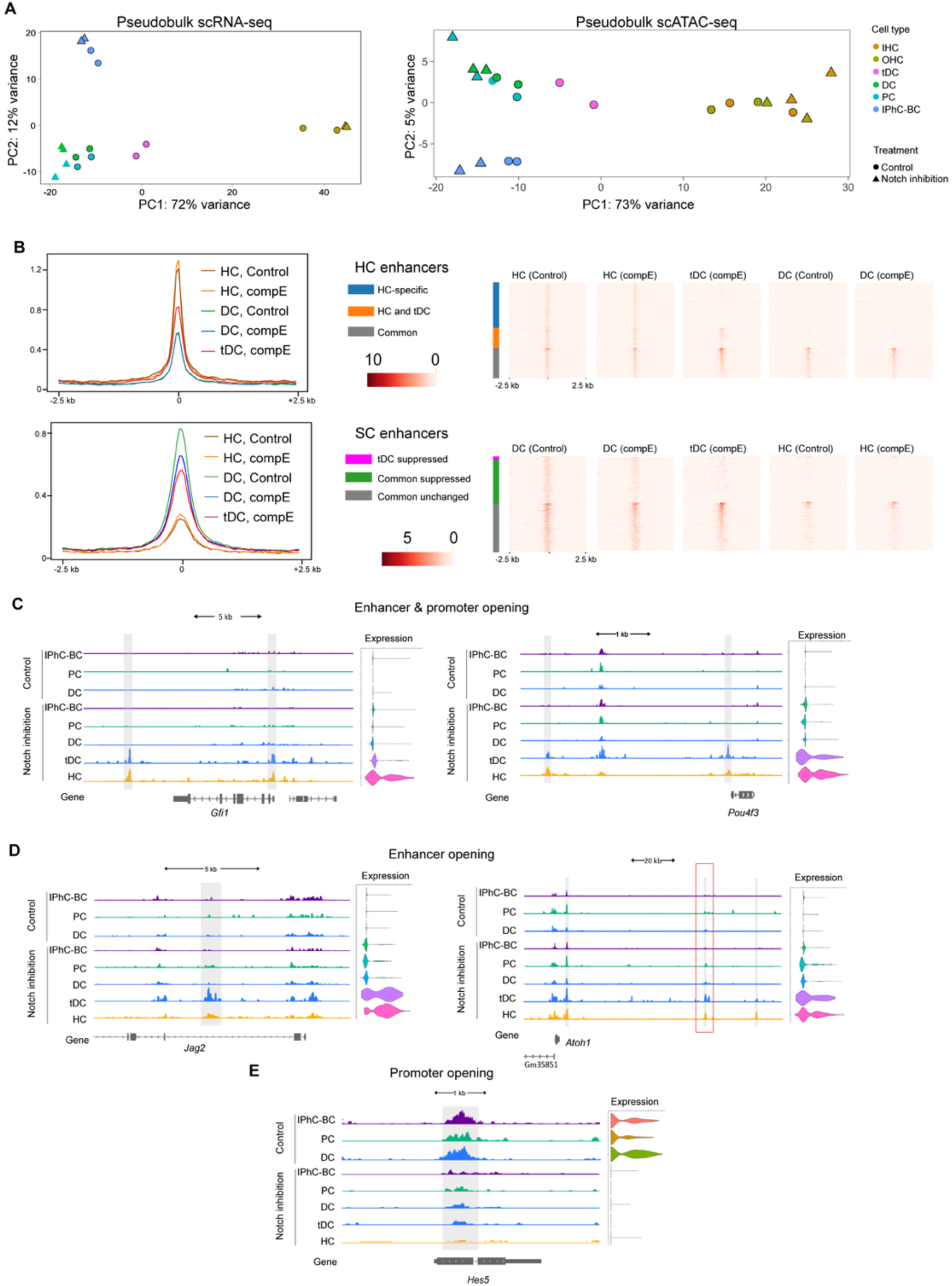
Coordinated transcriptional and epigenomic remodeling distinguishes tDCs from stable SCs. (**A**) Principal component analysis (PCA) of pseudobulk single-cell RNA-seq (left) and ATAC-seq (right) data showing clear separation of major cochlear cell types under control (DMSO) and Notch inhibition (compound E, compE) conditions, while in tDCs (pink) there was a clear shift with Notch inhibition, separating from stable DCs toward a HC like fate. (**B**) Average signal intensity profile and heatmaps of chromatin accessibility at enhancer categories (HC and SC enhancers). Top: HC enhancers are clustered into three groups - enhancers accessible exclusively in HCs, blue; enhancers accessible in both HCs and tDCs, orange; commonly accessible enhancers, gray. tDCs gain accessibility at a subset of HC-associated enhancers, closely resembling HCs under compound E treatment. Bottom: SC enhancers clustered into enhancers losing accessibility in tDC, magenta; enhancers with decrease in accessibility in all DCs, green; enhancers whose accessibility remains unaffected by Notch inhibition in grey. The majority of SC enhancers responding to Notch inhibition are affected in all DCs, but their accessibility loss is more prominent in tDCs. **(C-E)** Representative genome browser tracks illustrating distinct modes of regulatory remodeling with transcriptional changes following Notch inhibition (gray boxes highlight the locus changes). (**C**) enhancer + promoter gaining chromatin accessibility (*Gfi1*, *Pou4f3*); (D) enhancer-only increase in accessibility (*Jag2*, *Atoh1*); (E) decreasing promoter accessibility (*Hes5*); Violin plots indicate corresponding gene-expression levels. Data represent two independent biological replicates per condition.

Next, to identify regulatory elements underlying the transition, we analyzed chromatin accessibility at curated HC-specific and SC-specific enhancers (previously identified in (*16*)) (Fig. 4B and fig. S6A). Aggregate ATAC-seq profiles (metaplots, left) revealed that tDCs (red-compound E) exhibited elevated accessibility across HC enhancers, approaching levels observed in native HCs (brown/orange), whereas accessibility of HC enhancers in both control DCs (green) and DCs exposed to Notch inhibition but not converted (blue) remained low (Fig. 4B). Heatmaps confirmed this pattern and further revealed a subset of enhancers that were shared between tDCs and HCs but remained inaccessible in non-tDCs, indicating partial convergence toward an HC regulatory landscape.

Conversely, analysis of SC enhancers revealed selective loss of accessibility in tDCs (Fig. 4B and fig. S6A). While some SC enhancers were broadly reduced following Notch inhibition, a distinct subset (“tDC suppressed”, pink) was specifically reduced in tDCs, whereas stable DCs retained accessibility at these loci. We also found a group of SC enhancers that became closed in tDCs, matching the closed state seen in HCs (SC-tDC suppressed, green).

In contrast to distal regulatory elements, genome-wide metaprofiles revealed few changes in accessibility at either HC or SC gene promoters (previously identified by (*16*)) between compound E-treated and control (fig. S6B), reiterating the key role of enhancers in the establishment of cell-fate transcriptional programs. Within each cell type, including tDCs, promoter accessibility profiles overlapped closely between treatment conditions, indicating that the core promoter landscape remains largely conserved. However, a small but biologically meaningful subset of promoters displayed accessibility changes specifically in tDCs. These included key HC regulators such as *Pou4f3* (Fig. 4C), *Nhlh1*, *Neurod6*, and *Dll1* (table S4), which gained accessibility together with upregulation of their expression, while *Hes5* was the only SC-associated promoter that lost accessibility (Fig. 4E and table S4). In contrast, most promoters including HC (*Hes6*) and SC (*Heyl*) genes showed minimal differences across conditions (fig. S6C), suggesting that the core promoter landscape remains largely stable during this transition.

Notably, while *Atoh1* expression was modestly induced across several SC clusters, its chromatin accessibility remained largely unchanged except its enhancer 2 element, previously defined as an HC-specific regulatory region (*27*). This selective opening demonstrates that tDCs epigenetically engage the HC gene-regulatory network (Fig. 4D, red box). While *Pou4f3 and GFi1*, downstream targets of Atoh1, showed marked promoter accessibility gains and transcriptional induction specifically in tDCs (Fig. 4C), indicating that strong Atoh1 activation is required to unlock its regulatory elements. This pattern suggests a hierarchical sequence in which Atoh1, as an upstream master regulator, becomes transiently activated from an already accessible locus, subsequently inducing chromatin opening at downstream HC genes such as *Pou4f3*

Our locus-level analyses defined four regulatory modes underlying the transition from SC to HC identity: (1) coordinated enhancer + promoter activation *(Pou4f3, Gfi1*; Fig. 4C), (2) enhancer-specific priming (*Atoh1, Jag2*; Fig. 4D), (3) promoter-specific repression (*Hes5*; Fig. 4E), and (4) chromatin stability with transcriptional activation (*Hes6*) or chromatin stability with transcriptional silencing (*Hey1*) (fig. 5SC). This spectrum of remodeling underscores the stepwise and gene-specific nature of the regenerative response, in which only a subset of loci achieves full enhancer promoter activation.

**Fig. 5.**
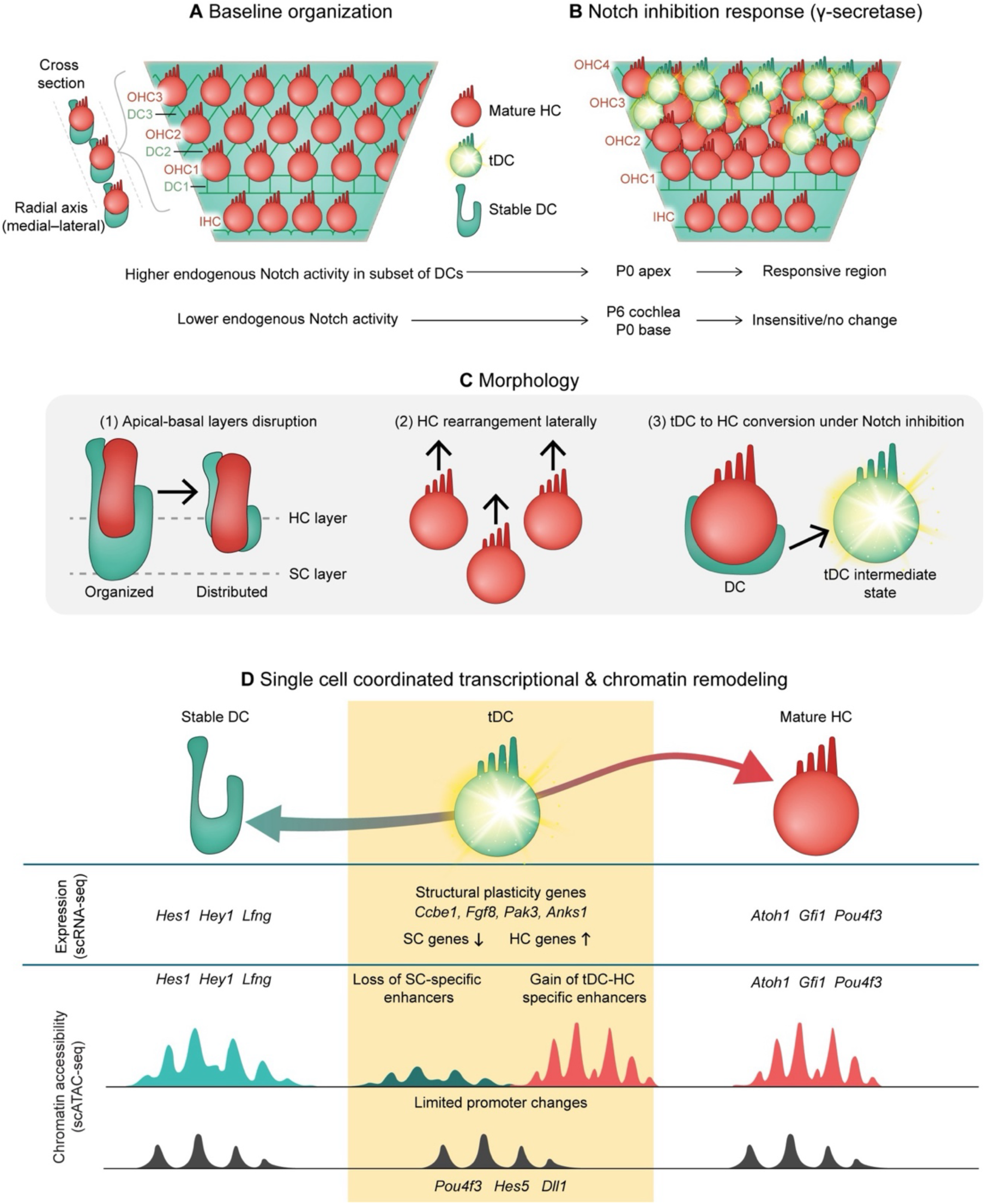
Integrated model summarizing the multilevel of tDCs to Notch inhibition in the neonatal cochlea. (**A**) Baseline organization of the neonatal (P0-1) organ of Corti. In the neonatal (P0-P1) organ of Corti, Deiters cells (DC1-DC3) form organized apical–basal layers beneath the three rows of OHC1-OHC3 and IHC. Baseline Notch activity is heterogeneous across DCs, and only a subset exhibits higher endogenous Notch signaling. Consistent with developmental restriction of regenerative competence, responsiveness is strongest at the P0 apex and reduced at the P0 base and by P6, correlating with lower endogenous Notch activity. (**B**) Notch inhibition response (γ-secretase). Following Notch inhibition, only a subset of DCs enters an intermediate transdifferentiating state (tDCs). (**C**) Morphological remodeling: Notch inhibition induces epithelial disorganization characterized by apical-basal disruption of DC layers and lateral rearrangement of pre-existing OHCs. Live imaging reveals that only a limited number of true DC-to-HC conversion events occur, whereas a substantial fraction of apparent supernumerary HCs arise from rearrangement and repositioning of pre-existing HCs within the epithelium. (**D**) Single-cell coordinated transcriptional and chromatin remodeling. At single-cell resolution, transition from stable DC to tDC and ultimately to mature HC is accompanied by coordinated changes in gene expression and chromatin accessibility. scRNA-seq: tDCs downregulate SC genes (*Hes1, Hey1, Lfng*) and upregulate HC regulators (*Atoh1, Gfi1, Pou4f3*), together with structural plasticity genes involved in adhesion and cytoskeletal remodeling (*Ccbe1, Fgf8, Pak3, Anks1*). scATAC-seq: Chromatin remodeling includes loss of SC-specific enhancer accessibility, gain of HC-specific enhancer accessibility, and selective promoter opening (e.g., *Pou4f3, Hes5, Dll1*), while many promoters remain stable.

While Notch inhibition broadly downregulated SC identity genes, most of these loci did not show corresponding changes in chromatin accessibility (fig. S5C). The only clear exception was *Hes5*, which lost promoter accessibility in all SCs including tDCs (Fig. 4E). This likely reflects the fact that many SC genes are controlled by constitutively open promoters and broadly accessible chromatin regions, allowing transcriptional repression to occur without major structural remodeling. In contrast, a subset of SC specific regulatory elements, particularly distal enhancers, showed clear loss of accessibility specifically in tDCs, indicating targeted selective removal of the SC enhancer landscape during fate conversion. This selective enhancer remodeling explains why only some SC genes show accessibility changes, while most are transcriptionally repressed without accompanying chromatin closure.

Motif analysis of enhancers shared between tDCs and HCs showed enrichment for Atoh1, Pou4f3, and Six1 motifs, indicating that tDCs open regulatory regions governed by core HC transcription factors (fig. 5SD and table S5).

Together, these findings demonstrate that tDCs selectively activate HC gene regulatory elements, while repressing SC associated regulatory regions, and that this regulatory rewiring occurs primarily at distal enhancers with limited but strategic promoter level remodeling at pivotal regulators that reinforce the transition toward an HC-like state.

## Discussion

Here we identify a fate-primed SC subpopulation in the neonatal cochlea. Using live imaging and single-cell multi-omics, we uncover how Notch inhibition selectively unmasks a rare subset of DC with transcriptional and epigenetic features that confer regenerative competence, while neighboring SCs remain refractory. Importantly, our data capture tDCs as an intermediate state, distinct from both stable SCs and mature HCs. This transitional identity is defined by enhancer remodeling and partial activation of HC gene programs, providing a snapshot of the earliest steps of fate conversion. These findings provide new insight into why Notch inhibition induces only limited regeneration and provide a mechanistic framework for differential SC plasticity.

Our study provides new insights into the cellular and molecular heterogeneity underlying the limited regenerative capacity, with a subtype dependent on the neonatal cochlea. Notch inhibition exhibited a limited regenerative potential in the neonatal stage, promoting OHC formation in the mid and apex turns by the transdifferentiation of DCs, mainly from the lateral row (DC3) (Fig. 1 and 2). This study establishes DCs as a uniquely responsive SC subtype and positions live imaging as a critical tool for resolving the interplay between molecular reprogramming and epithelial architecture during sensory regeneration. Beyond fate conversion, our live imaging uncovered morphological plasticity and migratory behaviors of SCs among Notch inhibition, including both responsive and non-responsive cells. This behavior disrupts the normal epithelial patterning, including the mixing of different epithelial layers. These dynamic cellular rearrangements indicate that Notch inhibition triggers coordinated changes in architecture alongside transcriptional reprogramming.

With the single-cell multi-omics, the picture was refined by uncovering significant heterogeneity in the regenerative competence of SCs. We identified that Notch inhibition does not elicit a uniform regenerative response across SCs, but instead, reveals distinct subpopulations with heterogenous plasticity and regenerative capacity (Fig. 3 and 4). Notably, we showed a subset of DCs that respond to Notch inhibition transcriptionally and epigenetically by transdifferentiating into HC, which we termed tDC, exhibiting dynamic morphological remodeling and activation of HC gene programs (*Atoh1*, *Pou4f*3, *Hes6*), increased chromatin accessibility at key HC loci such as *Pou4f3.* Strikingly, promoter accessibility was globally stable, selective promoter changes at key regulators (e.g., *Pou4f3*, *Dll1*, *Hes5*) reinforce the idea that transcriptional commitment is secured by a small number of pivotal nodes, whereas broad enhancer remodeling drives the wider fate competence of tDCs. Integration of accessibility profiles revealed distinct regulatory modes across the genome, including regions showing enhancer-specific, promoter-specific, or combined enhancer-promoter remodeling, while others remained unchanged (Fig. 4 and fig. S5B). Gene ontology analysis further supported this interpretation, highlighting enrichment for developmental and morphogenetic programs associated with adhesion, junctional remodeling, and epithelial differentiation, consistent with the cellular reorganization and structural plasticity observed in live imaging. In contrast, other SC subtypes, including IPhCs and BCs, remained transcriptionally silenced and epigenetically refractory to fate conversion despite effective Notch pathway repression, highlighting intrinsic differences in regenerative potential among SC populations. While live imaging demonstrated DC transdifferentiate toward OHC-like morphology, the single-cell multi-omic data did not clearly resolve an OHC fate. This may reflect the early time point analyzed, capturing cells in an intermediate transcriptional state before full OHC specification.

Our results thus extend existing models by providing a mechanistic framework for understanding differential SC plasticity. Rather than viewing Notch inhibition as a global regenerative cue, we propose a refined model in which the cochlear sensory epithelium comprises at least two functional SC classes: responsive SCs, which are molecularly and epigenetically primed for transdifferentiation, and "stable" SCs, which resist reprogramming even under permissive conditions, precisely identifying a fate-competent SC subpopulation. Transcriptional and epigenetic priming for HC fate was revealed using high-resolution live imaging and single-cell multi-omics, providing mechanistic insights into what distinguishes these responsive cells from non-responsive ones. These findings fill a critical gap in the field: while Notch inhibition has long been known to induce limited SC to HC conversion, the identity of the competent cells and the molecular basis of their selective responsiveness have remained unknown. By identifying this rare population at single-cell resolution, our study defines the molecular and epigenetic features that distinguish a fate-competent SC state. Since our multi-omic analysis captures a single early time point following Notch inhibition, it does not resolve the temporal sequence of events preceding entry into the transdifferentiating state. Future studies with finer temporal resolution will be required to determine whether competence emerges *de novo* upon pathway inhibition or reflects pre-existing heterogeneity within the DC population.

While our data identify tDCs at the transcriptional and epigenomic levels, the signals underlying their preferential emergence in the lateral (DC3) region remain to be determined. We note that future studies incorporating higher resolution signaling, transcriptional, and epigenetic spatial could provide additional insights into the regulatory states that predispose specific DC subtypes, such as DC3, to respond to Notch inhibition. Such approaches may clarify whether distinct signaling or chromatin priming underlies the selective responsiveness observed in DCs.

Our findings indicate that transcriptional repression of SC identity alone is insufficient to drive fate conversion; rather, only tDCs can overcome additional regulatory constraints to initiate the HC lineage program. The persistence of a transcriptionally silent state in stable DCs, PCs, and IPhCs-BCs despite robust Notch target downregulation suggests the existence of additional epigenetic barriers preventing HC gene activation. Thus, we provide the molecular definition of a fate-primed SC state and demonstrate how transcriptional and epigenetic priming combine to confer regenerative competence.

These findings suggest that low regenerative efficiency after Notch inhibition is due to intrinsic heterogeneity in epigenetic competence among SCs. Overcoming this bottleneck will require identifying the mechanisms that endow tDCs with their unique competence and, critically, how to induce this state in other SC types. This could involve targeted manipulation of signaling pathways (e.g., combined Notch and FGF inhibition) together with epigenetic reprogramming to unlock closed chromatin at HC regulatory loci. Moreover, the morphological plasticity and migratory behaviors we observed reveal that regeneration involves not just molecular reprogramming but also large-scale architectural remodeling that highlighting the importance of integrating tissue mechanics and patterning into regenerative approaches.

Future work should focus on elucidating the barriers that maintain stable SCs in a refractory state and on discovering strategies to convert them into responsive SCs. Approaches such as isolation tDC, *ex-vivo* organoids assay or CRISPR-based epigenetic activation of key HC-associated enhancers in stable SCs. These strategies could reveal the molecular switches that govern SC fate competence and provide a blueprint for reprogramming non-responsive cells. The concept of a fate-primed progenitor-like SC subpopulation may also extend beyond the cochlea to other tissues with limited regenerative capacity.

Our findings extend existing models by showing that regeneration in the cochlea involves both molecular and structural dimensions (Fig. 5). Notch inhibition uncovered coordinated transcriptional reprogramming, enhancer remodeling, and large-scale tissue dynamics, including rearrangement and mis-patterning. Together, these processes reveal that regenerative competence requires the integration of molecular priming (transcriptional and epigenetic) with architectural adaptability at the tissue level.

Together, these data support a model in which only a developmentally restricted subset of DCs undergoes coordinated transcriptional and epigenetic reprogramming, acquiring both HC fate determinants and structural plasticity required for epithelial reorganization.

In summary, this study demonstrates that regenerative competence in the neonatal cochlea is not uniformly distributed but confined to a distinct SC subpopulation with unique molecular, epigenetic, and morphological properties. This recognition of intrinsic heterogeneity and the identification of the fate-primed SC population provide a conceptual framework and practical roadmap for developing effective strategies to regenerate sensory HC in the damaged mammalian cochlea.

## Materials and methods

### Experimental animals

Experiments with mice were carried out at Tel Aviv University (TAU) and meets the guidelines of the NIH Guide for the Use of Laboratory Animals and is approved by the Institutional Animal Care and Use Committee (IACUC) of TAU, numbers TAU-MD-IL-2311-165-2 for Lfng-EGFP and TAU-LS-IL-2212-174-2 for Atoh1 mCherry. Animal usage for the single cell-multiome experiment was approved by IACUC 1187 at Creighton University.

Lfng (B6;FVB-Tg(Lfng-EGFP)HM340Gsat/Mmucd) mice, generated by the GENSAT project, were used to obtain labeled SC (*20, 28*). The mouse strain used was B6;FVB-Tg(Lfng-EGFP)HM340Gsat/Mmucd, RRID:MMRRC_015881-UCD, obtained from the Mutant Mouse Resource and Research Center (MMRRC) at the University of California at Davis, an NIH-funded strain repository. This strain was donated to the MMRRC by Nathaniel Heintz, Ph.D., from the Rockefeller University, GENSAT. Cryopreserved sperm were recovered to grow mice by the Cryopreservation Unit at the Gray Faculty of Medical and Health Sciences and were maintained on a C57BL/6 background. EGFP (Forward primer: CGA AGG CTA CGT CCA GGA GCG CAC; Reverse primer GCA CGG GGC CGT CGC CGA TGG GGG TGT, yielding a 300bp band. Math_Cherry_F - GGTAGTTTGCCGTAATGTGAG XFPN_R - CTCCTCGCCCTTGCTCACC, yielding 467 bp. Atoh1-mCherry mice were a gift to Prof. David Sprinzak from Prof. C Petit, Institut Pasteur, and were maintained on a C57BL/6 background. Genotyping was performed using the KAPA HS Mouse Genotyping Kit from Sigma (catalog no. KK7351 220).

### Cochlea culture

All cochleae were dissected from postnatal P0-1 and P6 from wild-type strain, Lfng-EGFP (*20*), and Atoh1 mCherry-Lfng-EGFP (*20, 25*) mice. Inner ears were isolated, placed on a dish with Dulbecco’s PBS w/o CA & MG Sartorius (catalog no. 02-023-1A) and cochlea were further dissected under the microscope and cultured for 24-48hr.

Cochleae were cultured with Cell-Tak or Millicell Standing Cell Culture 24 well (cat.no: PICM01250). Cochlear culture medium: (Gibco™ N-2 Supplement (100X) cat.no:17502048 Thermo Fisher Scientific, DMEM/F12 D6421-500ML Merck, Ciprofloxacin hydrochloride, enzo life sciences catalog number ALX-380-288G005 (1:1000) and with 400ul FBS- 500mL, Origin US SALE-SH30071.03IR25-40. The compound E 2uM, gamma secretase inhibitor from Abcam (AB142164-1-B) was dissolved in DMSO.

### Dissociation and FACS sorting

Cochlear tissues were pooled from four organs into a single tube and dissociated by adding 200 µl of Trypsin-EDTA solution (0.25%, T4049-100ML, Sigma) and PBS (1:1). The tube was placed in a water bath with gentle shaking at 300 rpm for 8 min. Following this, 100 µl of the dissociation solution was removed and replaced with 200 µl of PBS containing 10% FBS. A P200 pipette was then used to titrate the solution by pipetting it 300 times. The resulting cell suspension was passed through a 40 µm cell strainer to remove debris, followed by a 30-second centrifugation. Cells were then purified and collected using a BD FACSAria Flow Cytometer with a 100 µm nozzle. Flow cytometry analysis was performed at the Flow Cytometry Unit at the Research Infrastructure Core Facility at the Gray Faculty of Medical and Health Sciences, Tel Aviv University as a service. Standard flow cytometry files (.fcs) were compensated and analyzed using BD Biosciences software. Two populations were quantified: “single EGFP only” and “double-positive populations. The percentage of double-positive cells was calculated as: 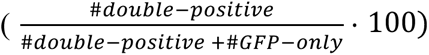. Quantification of responsive SC counts was based on events collected from the BD FACSAria. This approach controls for potential confounding factors such as dissociation efficiency, organ number, and cell viability.

### RNA sequencing and data analysis

SCs were sorted into RLT Plus lysis buffer from the Qiagen RNeasy Plus Micro Kit (Catalog no. 74034) and RNA was extracted according to the manufacturer’s instructions. The RNA quality was assessed using a TapeStation. Libraries were prepared from approximately 0.5-1 ng of total RNA from sorted cells (RIN > 7) using the SMART-Seq® v4 PLUS Kit (Takara Bio). During cDNA amplification, 10 PCR cycles were performed, and 16 PCR cycles were applied during library amplification. The prepared libraries were sequenced on the NextSeq™ 500 platform (Illumina) using the NextSeq 500/550 High Output Kit v2.5 (75 Cycles) (Illumina). Both library preparation and sequencing were performed at the Genomics Research Unit, Life Sciences Inter-Departmental Research Facility Unit, Tel Aviv University.

RNA-seq reads were pre-processed for quality and adaptor trimming using TrimGalore v0.6.0 with Cutadapt v1.15. STAR v 2.7.5c software was used to map the reads to GRCm38/mm10 mouse genome assembly and assign read counts to gene annotation.

Differential expression analysis was performed using DESeq2 v1.46.0. Genes with at least two-fold expression change and FDR<=0.01 were considered differentially expressed.

### Immunohistochemistry

Mice were euthanized by decapitation. The inner ears were carefully dissected under operating binoculars and fixed in 4% PFA for 2 h RT. After fixation, the inner ears were washed in PBS and stored at 4°C until further fine dissection. The organ of Corti was isolated in PBS under the operating binoculars. Specimens were permeabilized and blocked in 2% Triton X-100 and 10% normal goat serum for 2 h RT. Subsequently, the specimens were incubated overnight at 4°C with the appropriate primary antibody, diluted in Phosphate Green antibody diluent (Bar Naor Ltd.), following the manufacturer’s instructions. After washing, the specimens were incubated for 2 h RT with the relevant secondary antibody, diluted in PBS according to the manufacturer’s guidelines. Finally, the specimens were mounted in Fluorescent Mounting Medium (Aqueous) E18-E18 and imaged using a Leica SP8 confocal microscope.

### Live imaging

Cochlear explants were prepared and cultured as previously described (*25*). Briefly, Matrigel (Corning, cat. 356237) was diluted with phenol red-free DMEM/F-12 (Thermo Fisher Scientific, cat. no. 11039021) supplemented with HEPES and L-glutamine. Explants were positioned apical side down in a drop of diluted Matrigel on glass-bottom dishes and incubated at 37°C for 20 min to allow polymerization. To minimize tissue drift, a drop of agarose, low gelling temperature (Sigma-Aldrich, catalog number A4018) was placed on top, followed by 1 ml of culture medium containing either DMSO (vehicle control) or 2 µM compound E.

Time-lapse imaging was performed using a Zeiss LSM 880 confocal microscope (Zeiss, Oberkochen, Germany) equipped with a fast Airyscan detector, a temperature-controlled chamber (37°C), and a CO₂ regulator (5% CO₂). Images were acquired with Z-stacks of 40 optical sections at 0.7 µm intervals every 30 min over the course of the experiment. Acquisition was controlled with Zeiss ZEN software.

### Image analysis

Image processing and analysis were carried out in Fiji/ImageJ (NIH). Selected optical planes and time points were extracted from the raw datasets. For colocalization analysis, red and green channels were binarized, and colocalized signals were identified using the Image Calculator (logical AND operation). Double-positive cells at 48 h were manually identified and then tracked retrospectively to the initial time point.

Each tracked cell was assigned an initial DC row position by counting the number of SCs separating it from the outer OPC row. The final OHC row position was determined by the number of intervening HCs relative to the OPC row (Fig. 2C). Side views were generated using the “Reslice” function in ImageJ. Quantification of tracked cell distributions was visualized as bar plots generated in Python using Matplotlib. “Reslice” in ImageJ. Bar plot of tracked cells was created using Python matplotlib.

### Statistical analysis

All statistical analyses were conducted using Prism 10 (GraphPad Software, San Diego, CA). Specific statistical tests, and corresponding *P* values are reported in the figure legends. For multiple group comparisons, significance was adjusted using the Holm-Sidak post hoc method.

### Single-cell multi-omics, library prep and data analysis

Twenty-four h after compound E or DMSO treatment, cultured organs were collected and dissociated for single cell multi-omics analysis following the protocol by 10X Genomic Inc. Briefly, organs were collected into 1.5 ml tubes and digested with 200 µl 0.125% trypsin for 8 min at 37°C. Enzymatic digestion was stopped by adding 200 µl 10% FBS in DPBS, and then digested organs were triturated with a 200 µl pipette until no visible tissue chunks. The resulting single cell suspension was further processed following the Low Cell Input Nuclei Isolation protocol (10X Genomics, Nuclei Isolation for Single Cell Multiome ATAC + Gene Expression Sequencing, CG000365, Rev D). After counting the number of nuclei and morphological examination under a microscope, the nuclei suspension was processed by a 10X Chromium iX controller for RNA and ATAC libraries. Resulting sequencing libraries were sequenced on NovoSeq X Plus platforms with either 10B or 25B flowcells for approximately 200 million paired-end reads (150bp for read1/2, 10bp for index 1 and 24 bp for index 2) per libraries.

Raw reads were first trimmed to retain required bases (28+10+10+90bp for RNA and 50+8+24+49bp for ATAC) and then processed using cellranger-arc; and processed data were further analyzed using packages Seurat v4.0. Transcriptomic data were filtered, normalized, scaled, and then integrated for unsupervised cell clustering. Cell clusters were annotated based on known marker genes (*26*), and then differentially expressed genes between treated and control samples were identified using FindMarkers command from Seurat or DESeq2 after deriving pseudo-bulk data from scRNA (table S6). Based on the cell type annotation of scRNA, aligned ATAC reads were grouped by cell types to make pseudo-bulk ATAC data for downstream analysis.

Chromatin accessibility peaks were detected by MACS2 software (--nomodel --nolambda, otherwise with default parameters), using two pseudobulk ATAC-seq replicates of each sample as input. For downstream analyses, peaks from different samples were merged to produce a single ATAC-seq peak annotation, and reads coverage in each sample was calculated using bedtools coverage function. Peaks with differential accessibility were detected using DESeq2 v1.46.0. For visualization in genome browser, ATAC-seq reads coverage was calculated in 50-bp windows and normalized with bedtools coverage function.

De-novo motif enrichment and enrichment analysis of known transcription factor motifs in differentially accessible ATAC-seq peaks were performed by Homer software (table S5). Gene ontology annotation and GO terms enrichment analysis was performed using Metascape online platform.

## Acknowledgments

We thank the members of the Prof. Karen Avraham and Prof. David Sprinzak laboratories for valuable discussions and critical input on this work. We are grateful to Prof. Christine Petit and Dr. Raphael Etournay (Pasteur Institute, France) for providing the Atoh1-mCherry mouse line. We also thank Dr. Irena Shur (Flow Cytometry Core), Dr. Sasha Lichtenstein (Cellular and Molecular Imaging Center), and Dr. Rami Khosravi (Single Cell Genomics Core) at the Research Infrastructure Core Facilities (RICF) of the Gray Faculty of Medical and Health Sciences and Dr. Hila Kobo from the Rosalie and Harold Rae Brown Cancer Research Core Facility at the Faculty of Life Sciences, Tel Aviv University. This work was also supported by the Bellucci Translation Hearing Center and partially conducted within the Auditory & Vestibular Technologies Core and Innovative Genomics and Bioinformatics Core at Creighton University.

## Funding

Breakthrough Award, 2711/22 of the Israel Science Foundation (KBA)

Ernest and Bonnie Beutler Research Program of Excellence in Genomic Medicine (KBA)

Sagol Center for Regenerative Medicine, Tel Aviv University (DS, KBA)

## Supplementary Materials

**The PDF file includes:**

Figs. S1 to S6

**Other Supplementary Material for this manuscript includes the following:**

Movies S1 to S3

Tables S1 to S6

## Data and code availability

All sequencing datasets generated during this study are available in the NCBI Gene Expression Omnibus (GEO). Single-cell multi-omics data are available under accession number GSE310785, and bulk RNA-seq data are available under accession number GSE313875.

