## Supplementary Material for "Live imaging and multimodal profiling reveal transdifferentiation of a cochlear supporting cell subpopulation upon Notch inhibition"

Science Advances
AAAS

Supplementary Materials for

### Live imaging and multimodal profiling reveal transdifferentiation of a cochlear supporting cell subpopulation upon Notch inhibition

Lama Khalaily *et al.*

**The PDF file includes:**

Figs. S1 to S6

**Other Supplementary Materials for this manuscript include the following:**

Table S1
Tables S2 to S6 titles
Movies S1 to S3 titles

#### Supplementary Figures

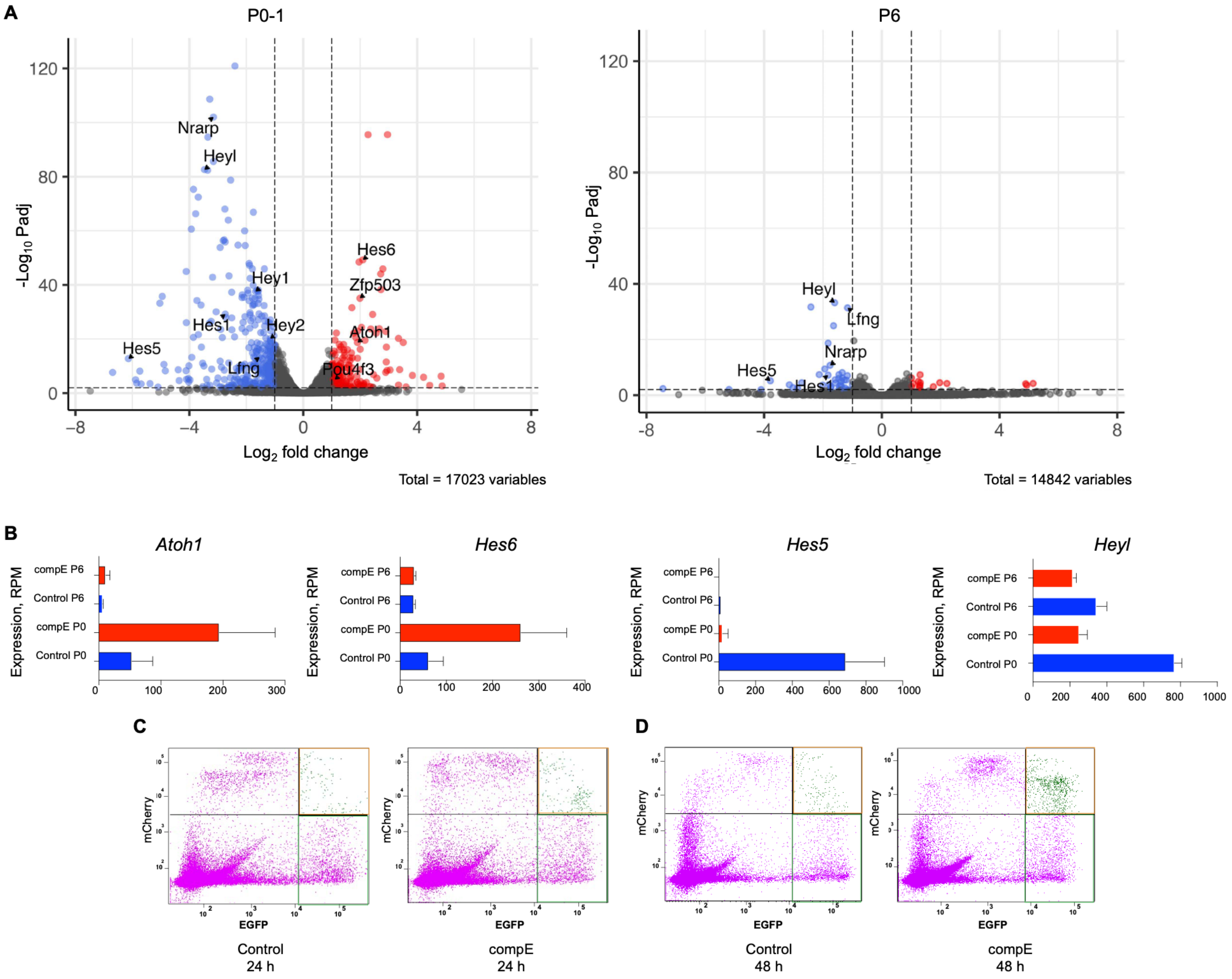


**Fig. S1. RNA-seq analysis validated stage-dependent changes in Notch target gene expression following Notch inhibition from supporting cells.** (**A**) Volcano plots showing differentially expressed genes upon Notch inhibition with compound E (compE) (2 μM) compared to DMSO control in cochlear samples at P0-1 (left panel) and P6-7 (right panel). At P0-1, 176 genes were significantly upregulated (red dots) and 346 genes were significantly downregulated (blue dots). In contrast, at P6-7, only 15 genes were significantly upregulated, and 40 genes were downregulated, indicating a reduced transcriptional response to Notch inhibition at older stages (n = 6-8 cochleae). (**B**) Expression of key hair cell (HC) and supporting cell (SC) markers across developmental stages and treatment conditions. HC markers *Atoh1 and Hes6* show increased expression upon Notch inhibition at P0-1, but this response is diminished at P6-7, correlating with restricted transdifferentiation potential at older stages. SC markers *Hey1 and Hes5* show a decrease in expression, indicating a shift toward HC fate. (**C, D**) Representative FACS profile for sorting of P0 Lfng$^{EGFP}$Atoh1$^{mCherry}$ cochleae after culture with DMSO or compound E for both 24 h and 48 h after culturing, used for sorting EGFP$^{+}$ only (green box) and EGFP$^{+}$mCherry$^{+}$ populations to quantify percentage of double positive cells (orange box). The double positive cells are found in both control and treatment Notch inhibition, but the number increased after compound E treatment, indicating the transdifferentiation events.

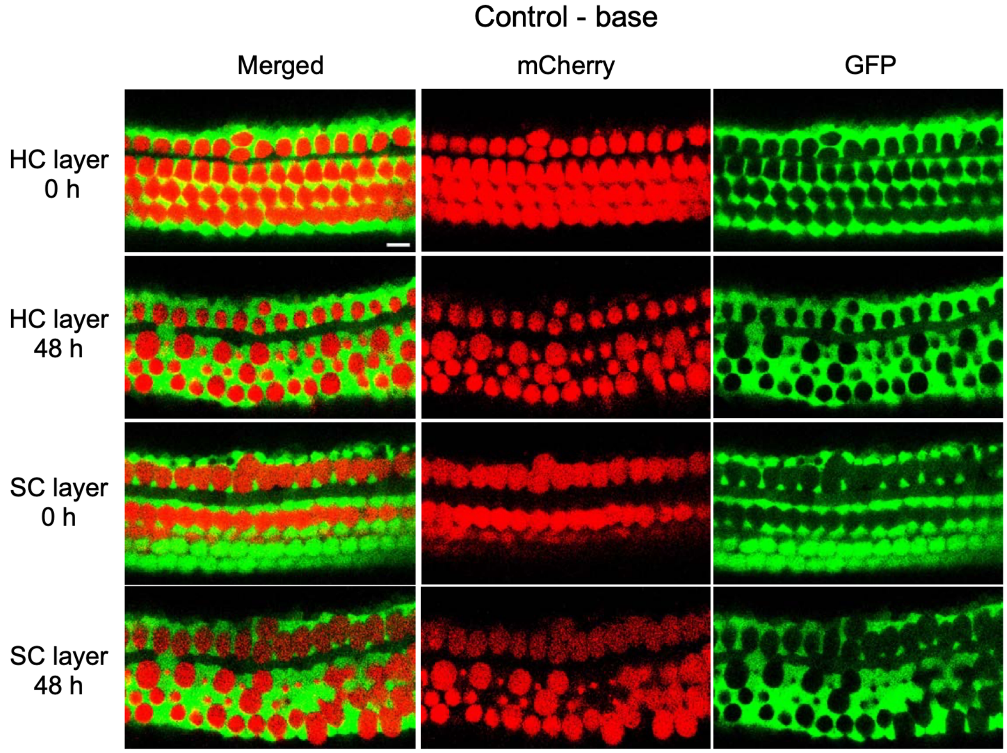


**Fig. S2. The precise pattern and layer organization in base treated with Notch inhibition.** Filmstrips from $Lfng^{EGFP}Atoh1^{mCherry}$ cochlear explants showing compound E-treated samples on base region. Images represent the HC and SC layers at 0 h and 48 h.

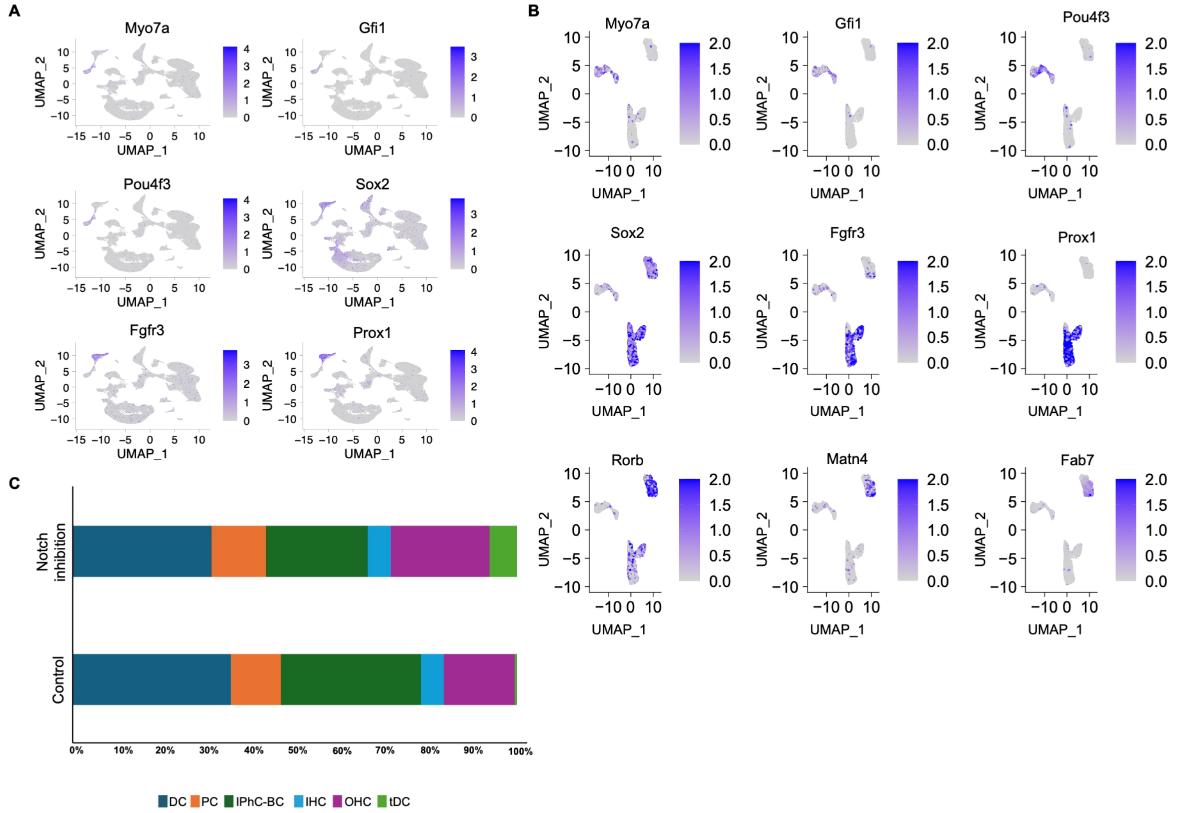


**Fig. S3. Marker gene expression and cluster distribution following Notch inhibition.** (**A**) Cluster annotation for expression of canonical marker genes across the UMAP projection used for identifying major cochlear cell populations. HC clusters were marked by *Myo7a*, *Gfi1*, and *Pou4f3*; DC and PC clusters by *Prox1* and *Fgfr3*; and IPhC-BC clusters by *Sox2*, consistent with their known molecular identities.(**B**) Marker gene projections onto the subcluster-level UMAP confirmed these assignments: tDCs expressed early HC-specific genes (*Myo7a*, *Pou4f3*, *Gfi1*), whereas stable DCs retained SC identity genes (*Sox2*, *Prox1*, *Fgfr3*). PCs expressed *Sox2*, *Fgfr3*, and *Prox1*, while IPhCs-BCs were characterized by *Sox2*, *Matn4*, *Fabp7*, and *Rorb*. (**C**) Proportions of cell per cluster in control versus Notch inhibition (compound E) samples shows enrichment of tDCs following Notch inhibition.

**A**

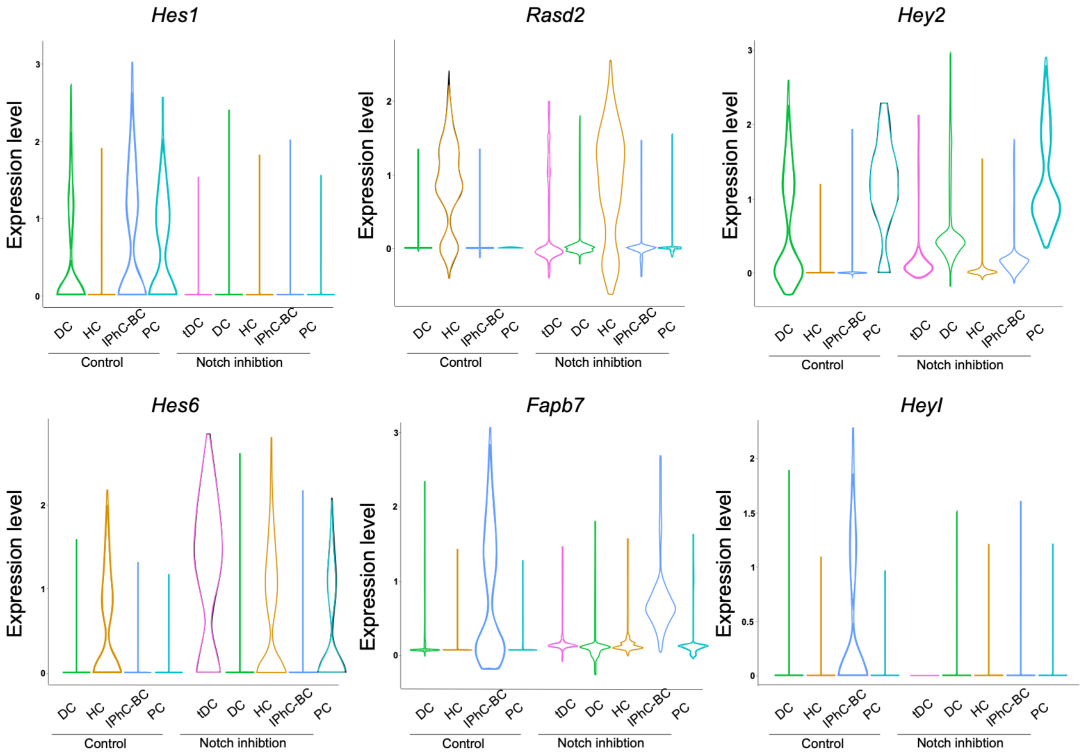


**B**

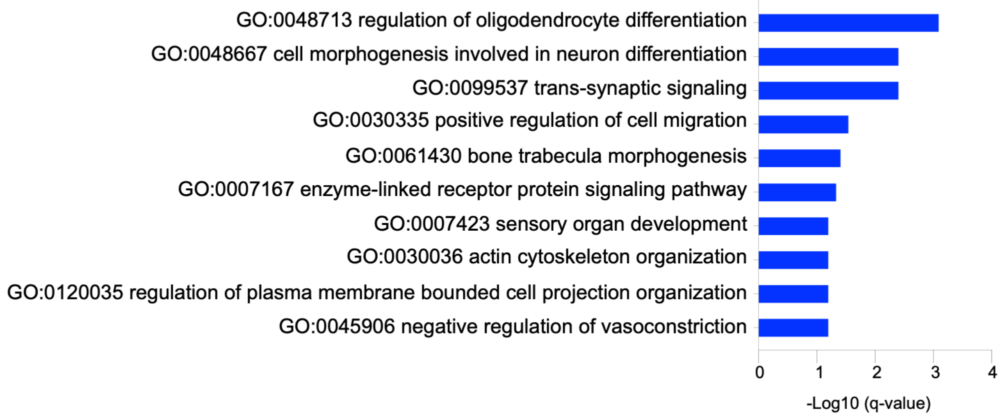


**Fig. S4. Gene expression changes and gene ontology for down-regulated genes.** (**A**) Gene expression changes shown by violin plots of canonical genes, *Hes1*, *Rasd1*, *Hey2*, *Hes6, Fabp7* and *Heyl*, and across cochlear cells from control to compound E-treated samples. Notch inhibition led to down regulation of *Hes1*, *Hey2*, *Fabp7* and *Heyl*, across all SC populations, while *Hes6* and *Rasd1* was selectively up regulated in tDCs, consistent with initiation of HC-associated transcriptional programs. *Hey2* expression remained high in PCs, and *Fabp7* persisted in IPhCs, indicating incomplete repression in these subtypes**.** (**B**) Gene Ontology enrichment analysis of down-regulated genes in tDCs identified processes related to cell morphogenesis, cytoskeletal reorganization, and migration, reflecting a transition from a stable epithelial to a motile, remodeling state.

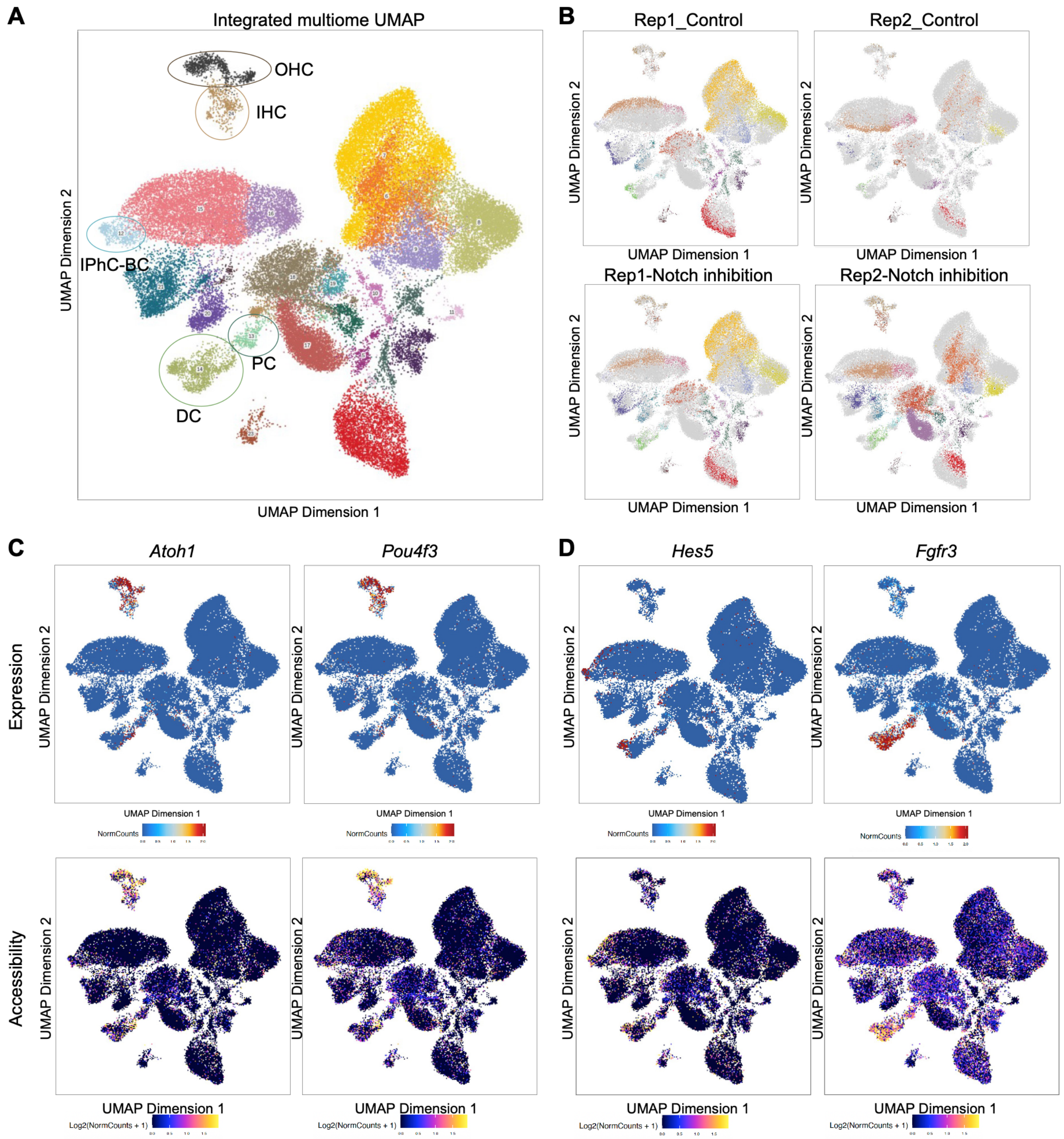


**Fig. S5. Multi-ome integration, sample origin, and concordance between gene expression and chromatin accessibility.** (**A**) UMAP projection generated from integrated sc-RNA and ATAC multi-ome data, colored by cluster identity and annotated with major cochlear cell populations. (**B**) The same UMAP coordinates colored by sample origin, showing cells from DMSO controls (top row) and compound E treatment (bottom row) across two biological replicates. (**C**) Representative feature plots showing concordance between gene expression (top) and chromatin accessibility (bottom) for HC-associated genes (*Atoh1, Pou4f3*). Accessibility patterns closely mirror transcriptional enrichment in HC clusters. (**D**) Feature plots of SC-associated genes. Both *Hes5*, a canonical Notch target, and *Fgfr3*, a SC marker, show enriched expression and accessibility within SC clusters, consistent with their established roles in SC identity.

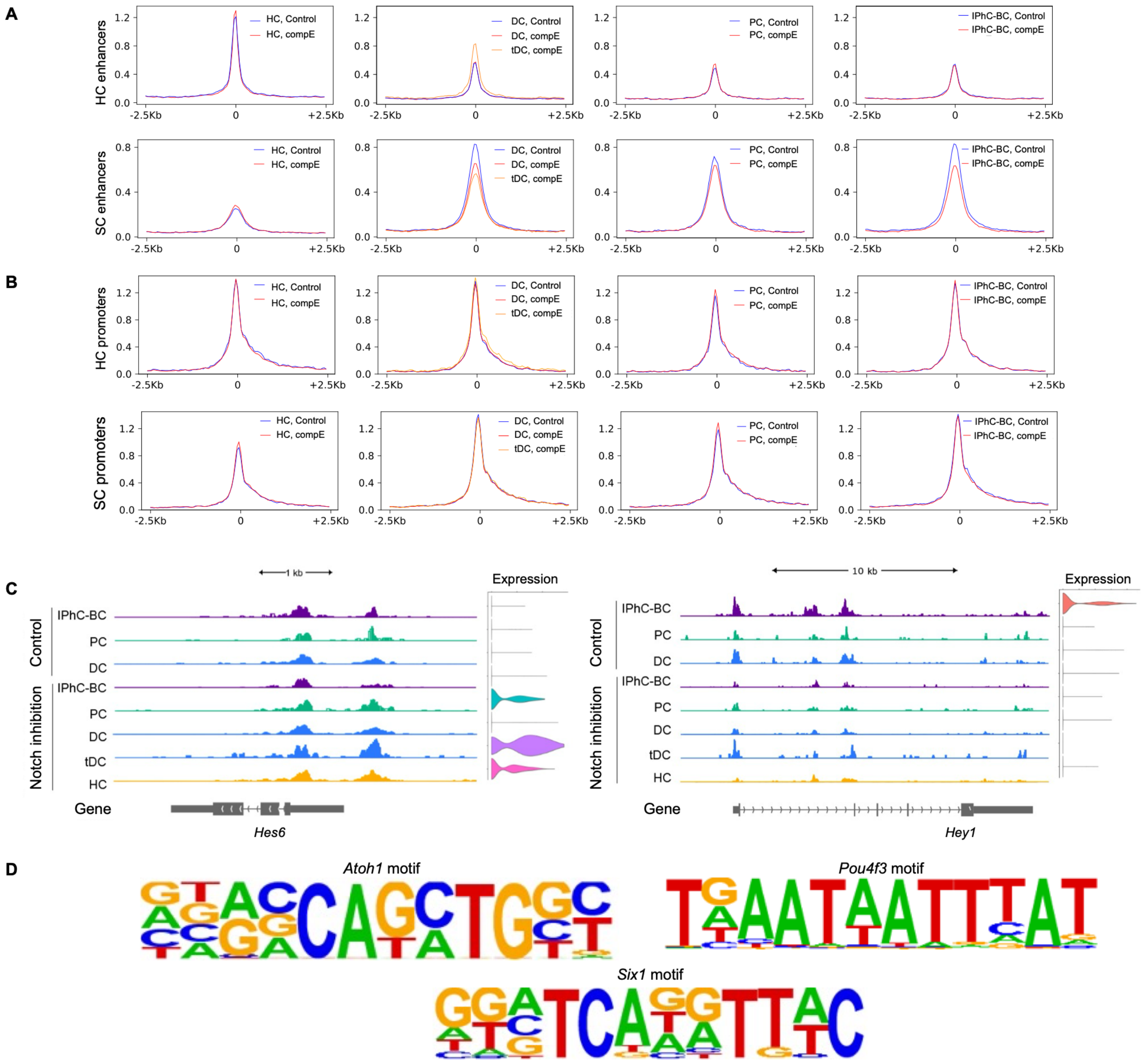


**Fig. S6. Chromatin accessibility remodeling and HC motif enrichment in tDCs following Notch inhibition.** (**A-B**) Aggregate accessibility plots for enhancer and promoter regions of HC and SC. (**A**) Accessibility profiles centered on transcription factor binding sites within HC-specific (top) and SC-specific (bottom) enhancers. tDCs showed a striking gain of accessibility at HC-specific enhancers compared with DCs and PCs, while SC enhancer accessibility showed some lower accessibility compared to control DCs**.** (**B**) Aggregate accessibility plots for promoter regions of HC and SC. Accessibility centered on transcription start sites of HC-specific (top) and SC-specific (bottom) promoters. Only a small subset of promoters displayed tDC-specific accessibility changes (summarized in table S4). (**C**) Representative ATAC-seq signal tracks showing accessibility profiles at *Hes6* and *Heyl* across cochlear cell types from control and **c**ompound E-treated samples. Each track represents aggregated accessibility from IPhC-BC, PC, DC, tDCs, and HC with their expression level. Notch inhibition maintained stable accessibility for *Hes6* and *Heyl* across all conditions. (**D**) Motif analysis of tDC-HC shared HC enhancers revealed significant enrichment for *Atoh1*, *Pou4f3*, and *Six1* motifs, indicating that tDCs engage canonical HC transcriptional regulators through enhancer activation.

**Supplementary Tables**

**Table S1. Overview of bulk RNA-seq datasets from P0 and P6 cochlear explants under Notch inhibition (compound E, compE) and control conditions (DMSO) (related to Fig. S1A).**

| Sample | Total reads, mln | Uniquely mapped, mln | Uniquely mapped, % |
|---|---|---|---|
| P0_compE_rep1 | 65.1 | 54.5 | 83.7 |
| P0_compE_rep2 | 26.9 | 22.0 | 81.7 |
| P0_compE_rep3 | 48.3 | 39.7 | 82.2 |
| P0_compE_rep4 | 44.4 | 36.6 | 82.4 |
| P0_DMSO_rep1 | 37.9 | 31.2 | 82.4 |
| P0_DMSO_rep2 | 54.6 | 45.1 | 82.6 |
| P0_DMSO_rep3 | 44.5 | 36.0 | 80.9 |
| P0_DMSO_rep4 | 42.0 | 34.5 | 82.0 |
| P6_compE_rep1 | 32.4 | 27.6 | 85.0 |
| P6_compE_rep2 | 35.4 | 30.9 | 87.3 |
| P6_compE_rep3 | 35.8 | 30.9 | 86.3 |
| P6_DMSO_rep1 | 30.4 | 25.9 | 85.2 |
| P6_DMSO_rep2 | 34.8 | 29.9 | 85.9 |
| P6_DMSO_rep3 | 54.4 | 45.9 | 84.4 |

**Table S2. Quantification and fate tracking of GFP$^+$mCherry$^+$ double-positive cells during live imaging in compound E-treated explants (related to Fig. 2).**

**Table S3. Gene Ontology enrichment (Biological Process) and annotation for up-regulated and down-regulated genes in tDCs compound E vs. DC-DMSO.**

**Table S4. Differential promoter accessibility analysis for tDCs compared to control DCs (DESeq2 results for promoter-associated peaks).**

**Table S5. Summary of motif enrichment analysis in differentially accessible regions (HOMER: de novo + known motifs).**

**Table S6. Pseudobulk RNA-seq differential expression analysis (DESeq2 results).**

**Supplementary Movies**

**Movie S1. *Lfng*$^{EGFP}$;*Atoh1*$^{mCherry}$ cochlear explant from the apex region treated with compound E, showing the emergence of GFP$^+$mCherry$^+$ double-positive cells of the OHC region**. Scale bar: 10 μm.

**Movie S2. *Lfng*$^{EGFP}$;*Atoh1*$^{mCherry}$ cochlear explant from the apex region treated with DMSO (control), showing stable tissue organization and absence of new mCherry$^+$ cells**. Scale bar: 10 μm.

**Movie S3. *Lfng*$^{EGFP}$;*Atoh1*$^{mCherry}$ cochlear explant from the basal region treated with compound E, showing no detectable appearance of GFP$^+$mCherry$^+$ double-positive cells, indicating minimal responsiveness to Notch inhibition at the base**. Scale bar: 10 μm.
